# Neurochemical signaling of reward and aversion by ventral tegmental area glutamate neurons

**DOI:** 10.1101/2020.05.18.103234

**Authors:** Dillon J. McGovern, Abigail M. Polter, David H. Root

## Abstract

Ventral tegmental area (VTA) glutamate neurons signal and participate in reward and aversion-based behaviors. However, the neurochemical mechanisms that underlie how these neurons participate in diverse motivated behaviors is unknown. We used a combination of optical sensors to identify how distinct neurochemical inputs to VTA glutamate neurons participate in motivated behavior. Glutamate inputs to VTA glutamate neurons increased for both reward- and aversion-predicting cues and aversive outcomes, but subpopulations of glutamate inputs were increased or decreased by reward. For all cues and outcomes, GABA inputs to VTA glutamate neurons decreased and GCaMP-measured neuronal activity increased. GCaMP recordings also showed that VTA glutamate neuronal activity discriminated between the omission and receipt of an expected reward, but glutamate and GABA inputs to these neurons did not. Electrophysiological recordings in coordination with our sensor data suggest that glutamate inputs, but not GABA inputs, principally regulate VTA glutamate neuron participation in diverse motivated behaviors.

## Introduction

The Ventral Tegmental Area (VTA) is a cellularly heterogeneous mid-brain structure^1–4^ that plays important roles in reinforcement, reward^5^, aversion^6–8^, and drug-seeking behavior^9^. Dopamine neurons predominate the VTA and contribute to reinforcing behavioral action via mesocorticolimbic targets^10^. Additionally, local VTA GABAergic neurons may disinhibit VTA dopamine neurons to support reward^11–13^ or may disrupt reward processing and promote aversion following their activation^14,15^. More recently a distinct subpopulation of VTA neurons that expresses the vesicular glutamate transporter (VGluT2) has been identified in the medial VTA of mice, rats, non-human primates, and humans^2^. These neurons release glutamate both locally within the VTA and distally to reward and aversion-related brain structures, such as lateral habenula and nucleus accumbens^1^. Initial exploration has revealed that VTA VGluT2 neurons heterogeneously participate in reward and aversion-based behaviors.

Photo-tagged VTA VGluT2 neuronal recordings show that nearly all of these cells increase their firing to aversive facial air puffs^16^. Interestingly, firing within these neurons is heterogenous for rewarding stimuli. Subsets of VTA VGluT2 neurons are reliably distinguished by whether they increase or decrease firing following reward receipt^16^. This heterogeneity in firing likely contributes to the functional diversity of these neurons in motivated behavior. Optogenetic stimulation of VTA VGluT2 neurons can result in either place preference or place aversion^17–19,20^. Stimulation parameters and projection targets both influence when mice exhibit reward or aversion-based behavior following VTA VGluT2 neuron stimulation, which indicates that the role of these neurons in motivation is complex. Our work in the following experiments aimed to determine the neurochemical mechanisms that underlie how VTA VGluT2 neurons signal reward and aversion. To accomplish this, we recorded from mice expressing genetically-encoded neurotransmitter sensing fluorescent reporters (SnFRs) to capture glutamate and GABA input to VTA VGluT2 neurons. We found that glutamate inputs to VTA VGluT2 neurons were divided into two populations, one population that increases for reward and another that decreases. In contrast, glutamate inputs homogeneously increased in response to aversive stimuli. GABA inputs to VTA VGluT2 neurons reliably decreased following both rewarding and aversive stimuli. While GABA inputs were altered by motivation-related stimuli and behaviors, our electrophysiological recordings indicated that glutamate input primarily regulates the spontaneous firing of these neurons. Together, these results 1) provide new neurochemical insights into how VTA VGluT2 neurons integrate multiple neurotransmitters during reward and aversion, and 2) suggest that glutamate is the primary regulator of VTA VGluT2 neuron activity and contributes to the diversity in signaling of reward and aversion by these neurons.

## Materials and Methods

All animal procedures were performed in accordance with National Institutes of Health Guidelines, and approved by the University of Colorado Institutional Animal Care and Use Committee or the George Washington University Institutional Animal Care and Use Committee.

### Intracranial injections and fiber optic implants

Male and female VGluT2-IRES::Cre mice (Slc17a6^tm2(cre)Lowl/J^; Jackson Laboratories, RRID:IMSR_ JAX:016963) (18-40g) were used (n=42). Mice were anesthetized with 1-3% isoflurane. AAV2/1-hSyn-FLEX-SF-iGluSnFR-A184S (n=11, 8 male, 3 female, titre 2×10^12^), AAV2/1-hSyn-FLEX-iGABASnFR-F102G (n=13, 11 male, 2 female, titre 2×10^12^), AAV1-hSyn-FLEX-GCAMP6m (n=7, 3 male, 4 female, titre 2×10^12^), or AAV8-EF1α-DIO-eYFP (n=11, 7 male, 4 female, titre 3×10^12^) was injected in VTA (500 nl; 100 nl/min; −3.2 mm anteroposterior, 0.0 mm mediolateral, −4.4 mm dorsoventral) using an UltraMicroPump, Nanofil syringes, and 35 gauge needles (WPI). Syringes were left in place for 10 min following injections to minimize diffusion. For neurotransmitter and calcium imaging experiments, an optic fiber (400 μm core diameter, 0.66 NA, Doric Lenses) was implanted dorsal to VTA (−3.2 anteroposterior, −1.0 mediolateral at 9.5, −4.2 dorso-ventral) and secured with skull screws and dental cement. No statistically significant differences between male and female mice were observed; all within-sensor data were therefore pooled.

### Reward task behavioral training

After at least three weeks following injection of viral constructs encoding iGluSnFR, iGABASnFR, or GCaMP, mice were food restricted to 85% free-feeding body weight. Mice were brought to chambers equipped with a three-dimensionally printed reward receptacle coupled to a syringe pump, two nose poke devices with cue lights, and a speaker (Med-Associates). Mice were initially trained in the reward task. During the reward task, either a conditioned stimulus (6 kHz tone and cue light activation for 10 sec; RewCS+) was presented that co-terminated with the delivery of 8% sucrose to the reward receptacle (20 μl) or a neutral conditioned stimulus (white noise for 10 sec; RewCS-) was presented that resulted in no sucrose delivery. RewCS+ and RewCS- trials were randomly presented, separated by a 60-120 sec variable inter-trial interval. Each training session ended after 30 RewCS+ and 30 RewCS- trials. The percentage of trials in which mice entered the reward port during RewCS+ or RewCS- trials was collected daily. Once mice reached at least three consecutive days of 70% or greater RewCS+ trials entering the reward port, neuronal recording commenced. During the neuronal recording, 40 RewCS+ and 40 RewCS- trials were presented. In addition, 90% of RewCS+ trials resulted in reward delivery and 10% of RewCS+ trials resulted in reward omission (termed reward CS+ error trials, Re-wCSe). One iGABASnFR mouse did not meet reward recording criterion and was only neuronally examined in the aversion task.

### Aversion task behavioral training

After recording neuronal activity in the reward task, mice were fed *ad libitum*. Mice were returned to the conditioning chamber over four days. The chamber context was altered by removing the reward receptacle, and cue lights remained illuminated for the entire duration of the task. During the aversion task, either a conditioned stimulus (10 kHz tone, 10 sec; ShkCS+) was presented that co-terminated with the delivery of foot shock (0.5 sec, 0.5 mA) or a neutral conditioned stimulus (white noise for 10 sec; ShkCS-) was presented that resulted in no foot shock. ShkCS+ and ShkCS- trials were randomly presented, separated by a 60-120 sec variable inter-trial interval. Each training session ended after 10 ShkCS+ and 10 ShkCS- trials. After three days of training, neuronal recording commenced. A separate group of wild type C57BL/6J mice (n=7 male) received the same training on the aversion task as described. However, prior to foot shock conditioning, the ShkCS+ and ShkCS-cues were randomly presented in the manner described above except that foot shock was not delivered. Four days of foot shock conditioning with the ShkCS+ and ShkCS-then commenced as described above. Video recordings (10Hz, Synapse, TDT, RRID:SCR_006495) were collected on the first session without foot shock delivery and during the last session with foot shock delivery. Total seconds of freezing, defined as the absence of movement except for breathing, was scored from videos during the ShkCS+ and ShkCS-cues.

### Neurotransmitter and calcium fiber photometry recordings

iGluSnFR, iG-ABASnFR, or GCaMP6m was excited at two wavelengths, 465nm and 405 nm isosbestic control, by amplitude modulated signals from two light-emitting diodes reflected off dichroic mirrors and then coupled into an optic fiber as previously described^21^. Sensor signals emitted and their isosbestic control emissions were returned through the same optic fiber and acquired using a femtowatt photoreceiver (Newport), digitized at 1kHz, and then recorded by a real-time signal processor (TDT). Behavioral timestamps of RewCS+ onset, RewCS-onset, RewCSe onset, reward port entries, reward delivery, ShkCS+ onset, ShkCS-onset, and foot shock delivery were digitized in Synapse software (TDT, RRID:SCR_006495) by TTL input from Med-PC (RRID:SCR_012156).

### Neurotransmitter and calcium signal analyses

Analysis of the recorded neurotransmitter and calcium signals was performed using custom-written MATLAB scripts (RRID:SCR_001622). Signals (465nm and 405nm) were down sampled (10X) and peri-event time histograms were created trial-by-trial between −20 sec and +30 sec surrounding each event analyzed. For each trial, data was detrended by regressing the isosbestic control signal (405nm) on the sensor signal (465nm) and then generating a predicted 405nm signal using the linear model generated during the regression. The predicted 405nm channel was subtracted from the 465nm signal to remove movement, photo-bleaching, and fiber bending artifacts (dF)^21^. Normalized dF was calculated by z-scoring each trial, where the mean and standard deviation of the z-score was calculated between −3 to 0 sec.

### Slice preparation

Male and female VGluT2-IRES::Cre mice (N=10, 5 male and 5 female) were injected with Cre-dependent AAV8-EF1α-DIO-eYFP at The University of Colorado. After at least three weeks, mice were shipped to George Washington University for slice physiology recording. After one week of acclimation, acute slices were prepared as previously described^22^ from deeply anesthetized mice. Mice were perfused with 34°C NMDG ringer (in mM): 92 NMDG, 2.5 KCl, 1.2 NaH_2_PO_4_, 30 NaHCO_3_, 20 HEPES, 25 glucose, 5 sodium ascorbate, 2 thiourea, 3 sodium pyruvate, 10 MgSO_4_, 0.5 CaCl_2_^23^. Following perfusion, the brain was rapidly dissected and horizontal slices (220 μM) were prepared in warmed NMDG ringer using a vibratome. Slices recovered for 1 h at 34°C in oxygenated HEPES holding solution (in mM): 86 NaCl, 2.5 KCl, 1.2 NaH_2_PO_4_, 35 NaHCO_3_, 20 HEPES, 25 glucose, 5 sodium ascorbate, 2 thiourea, 3 sodium pyruvate, 1 MgSO_4_, 2 CaCl_2_^23^ and then were held in the same solution at room temperature until use.

### Electrophysiology

Electrophysiological recordings were performed using a Sutterpatch integrated patch amplifier. Midbrain slices were continuously perfused at 1.5–2 mL/min with ACSF (28–32°C) containing (in mM): 126 NaCl, 21.4 NaHCO_3_, 2.5 KCl, 1.2 NaH_2_PO_4_, 2.4 CaCl_2_, 1.0 MgSO_4_, 11.1 glucose. VTA VGluT2 neurons were identified by eYFP fluorescence. For whole-cell recordings, borosilicate glass patch pipettes (3-5 MΩ) were filled with (in mM): 135 K-gluconate, 10 HEPES, 4 KCl, 4 ATP-Mg, and 0.3 GTP-Na. Whole-cell firing was measured in response to current injections at baseline and after ten minutes of drug wash-in. Cell attached recordings were obtained in the loose-patch configuration^24^. For cell-attached recordings, pipettes were filled with (in mM): 150 NaCl, 10 HEPES, 10 Glucose, 2.5 CaCl_2_, 1.3 MgSO_4_, 3.5 KCl. Cell attached firing rates were determined over a one-minute period at the end of a five-minute baseline or a one-minute period at the end of a 10-minute drug wash-in. DNQX (10 μM), APV (50 μM), isoguvacine (10 μM), AMPA (1 μM) and NMDA (10 μM), and bicuculline (10 μM) were bath applied. All recordings were performed in the presence of 1 μM strychnine to block glycine receptors. All salts and strychnine hydrochloride were purchased from Sigma-Aldrich (St. Louis, MO). DNQX, APV, isoguvacine, NMDA, and bicuculline were purchased from Tocris Biosciences.

### Statistical analyses

Statistical analyses were performed in SPSS (IBM, RRID:SCR_002865) or Graphpad Prism (RRID:SCR_002798). For the reward task, the average percent of RewCS+ and RewCS- trials that mice entered the reward port during the first three days of training and last three days of training was analyzed for each sensor group using repeated measures ANOVA. For the aversion task, the average number of seconds freezing in response to the ShkCS+ and ShkCS-during the first session with no foot shock conditioning and the final session with foot shock conditioning was analyzed using repeated measures ANOVA. For both tasks, Sidak pairwise comparisons followed up significant main effects or interactions.

For neurotransmitter sensor and calcium imaging, normalized dF (z-score) of neural activity was analyzed separately for each sensor (iGluSnFR, iGABASnFR, GCaMP6m). Activity −6 to −3 seconds prior to each cue was used to establish a baseline for their forthcoming cue and outcome^25^. Cue-related activity was examined 0 to 2 seconds following cue onset. Reward-related activity was examined following reward onset, 8.5 to 10 seconds following the RewCS+, and this timing was applied for non-reward activity following the RewCS-as well as for error-related activity following the RewCSe cues. Shock-related activity was examined following shock onset, 9.5 to 11.5 seconds following the ShkCS+, and this timing was applied for the non-shock activity following the ShkCS-. iGABASnFR signaling uniformly decreased from baseline for all cues and outcomes. Therefore, the minimum baseline value during the baseline epoch was compared with the minimum cue value during the cue epoch, and with the minimum outcome value during the outcome epoch (reward, shock, etc). GCaMP signaling uniformly increased for all cues and outcomes. Therefore, the maximum baseline value was compared with the maximum cue and outcome values. iGluSnFR signaling was increased by reward-related cues but either increased or decreased at reward. Due to the different patterns in reward-related activity within the iGluSnFR group, cue-related increases in activity were examined separately from reward-related activity. To examine cue-related activity, the maximum baseline value was compared with the maximum cue value using a paired t-test. We next compared the reward-related activity separately in reward-decreasing and reward-increasing iGluSnFR mice. The reward-increasing iGluSnFR subgroup compared the maximum baseline value with the maximum reward value using a paired t-test. The reward-decreasing iGluSnFR subgroup compared the minimum baseline value with the minimum reward value using a paired t-test. Unless otherwise stated, all analyses of neuronal activity first consisted of repeated measures ANOVA examining differences between baseline, cue, and outcome. If the assumption of sphericity was not met (Mauchley’s test), the Greenhouse-Geisser correction was used. Simple contrast tests followed up significant main effects of epoch examining the difference between baseline and cue, or baseline and outcome.

Electrophysiological data were analyzed using Sutterpatch software and Graphpad Prism. Cell-attached recordings of percent spontaneous firing rate from baseline in the presence of DNQX/APV, bicuculline, AMPA/ NMDA, and isoguvacine were analyzed using one sample t-tests. Current clamp recordings of the number of action potentials under baseline and DNQX/APV/bicuculline conditions were analyzed using repeated measures ANOVA.

### Data Availability

All custom MATLAB photometry scripts are publicly available at https://www.root-lab.org/code. All other relevant data are available from the corresponding author on reasonable request.

## Results

### Pavlovian Reward Task

Behavioral and neural data was collected from VGluT2::Cre mice expressing Cre-dependent iGABASnFR (to detect GABA release onto VTA VGluT2 neurons), iGluSnFR (to detect glutamate release to VTA VGluT2 neurons), or GCaMP6m (to detect changes in intracellular calcium of VTA VGluT2 neurons) using fiber photometry (**Supplemental Fig. 1**). Mice in each cohort initially trained on a Pavlovian reward paradigm where the termination of a RewCS+ resulted in the delivery of sucrose reward. Mice were concurrently trained to associate a neutral cue (RewCS-) with the absence of reward. Prior experimentation established that approximately half of VTA VGluT2 neurons change firing rates when an expected reward is omitted, suggesting VTA VGluT2 neurons can be sensitive to reward prediction error^16^. To identify whether glutamate or GABA inputs to VTA VGluT2 neurons signal prediction error, 10% of RewCS+ trials resulted in the omission of reward, termed error trials (RewCSe). After each mouse reached the established behavioral criterion (see *Methods*) they were recorded fiber photometrically.

### GABA inputs to VTA VGluT2 Neurons during Pavlovian Reward

Over the course of training, iGABASnFR mice learned to increase reward-port entries depending on the cue presented, F(1,18) = 14.5, p < 0.001. Reward port entries significantly increased in response to the RewCS+ on the last 3 days of training compared with the first 3 days of training, p < 0.0001. There were no statistically significant changes in reward port entries for RewCS-cues over training, p = 0.68 (**Fig. 1c**). iGABASnFR activity, which captures GABAergic transmission to VTA VGluT2 neurons, was significantly modulated by reward trials, F(2,22) = 60.02, p < 0.001. iGABASnFR activity was significantly decreased at the RewCS+ cue, F(1,11) = 5.65, p < 0.05, and GABA signaling further decreased at reward presentation, F(1,11) = 97.57, p < 0.001, when compared to baseline (**Fig. 1d-f**). iGABASnFR activity was also significantly modulated by reward omission trials, F(2,22) = 21.29, p < 0.001. During RewCSe trials when expected sucrose reward was omitted, iGABASnFR activity significantly decreased at the time of the expected reward, F(1,11) = 37.14, p < 0.001, suggesting that GABA input to VTA VGluT2 signals reward anticipation and are not sensitive to reward omission (**Fig. 1g-i**). For RewCS- trials that predicted no reward outcome, iGABASnFR activity was not significantly different from baseline, F(2,22) = 2.650, p = 0.09 (**Fig. 1j-l**).

**Figure 1.**
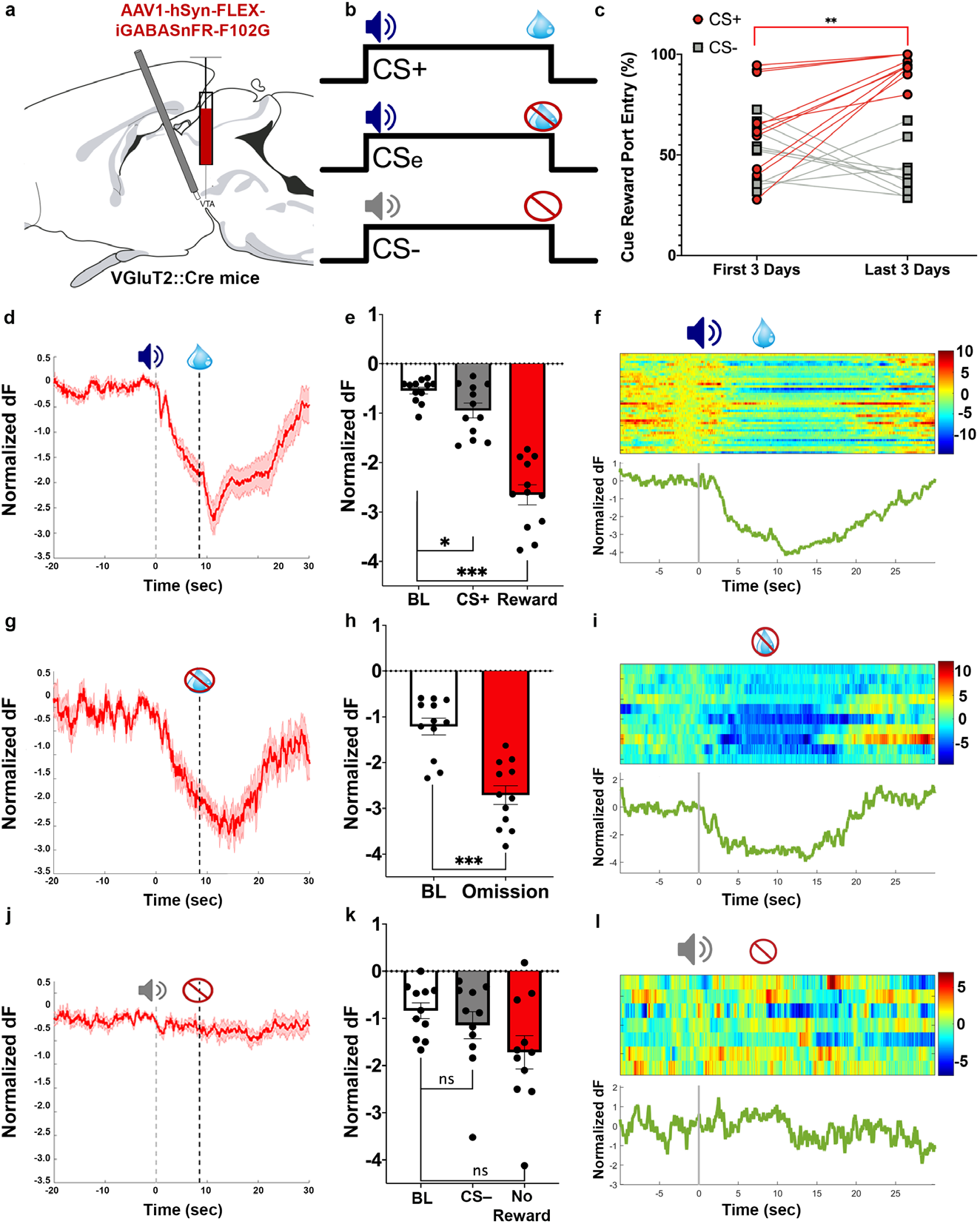
GABA input (iGABASnFR) to VTA VGluT2 neurons during Pavlovian Reward. a) VGluT2:: Cre mice were injected in the VTA with an AAV encoding Cre-dependent iGABASnFR and an optic fiber was implanted in the VTA. b) Pavlovian Reward Task. During training sucrose delivery was paired with CS+ tone but not CS- tone. During the recording, CSerror trials sounded the CS+ and resulted in no sucrose delivery (CSe). c) Mice learned to discriminate between reward-predictive cues (CS+, red) and neutral cues (CS-, grey). d) Normalized z-score (dF) for all iGABASnFR mice over time (group mean: solid line, S.E.M: shading). Decrease in GABA signaling at CS+ and sucrose presentation. e) Normalized dF of CS+ and reward compared to baseline. f) Sample trial-by-trial heat map and average trace from a representative iGABASnFR animal across reward trials. g) Normalized dF for all iGABASnFR mice during reward error trials (group mean: solid line, S.E.M: shading). GABA input decreased from baseline at reward omission h) Normalized dF of reward omission compared to baseline. i) Sample trial-by-trial heat map and average trace from a representative iGABASnFR mouse across error trials. j) Normalized dF for all iGABASnFR mice over time for neutral trials (CS-; group mean: solid line, S.E.M: shading). k) Normalized dF of CS- and non-reward compared to baseline. l) Sample trial-by-trial heat map and average trace from a representative iGABASnFR mouse across neutral CS- trials that resulted in reward-port entry. * p < 0.05, ** p < 0.01, *** p < 0.001.

### Glutamate inputs to VTA VGluT2 Neurons During Pavlovian Reward

iGluSnFR mice learned to emit head entries into the reward port over training depending on the cue presented, F(1,22) = 34.11, p < 0.0001 (**Fig. 2a**). Mice significantly increased reward port entries in response to the RewCS+ between the first and last three days of training, p < 0.0001, indicating that mice learned to increase responding for reward trials (Fig 2b). RewCS-head entries did not change over training, p = 0.99. iGluSnFR activity significantly increased following RewCS+ cues, t(11) = 2.27, p < 0.05. However, in contrast to the uniform decreased iGABASnFR activity at reward, subsets of iGluSnFR sensor mice showed divergent iGluSnFR activity at reward (**Fig. 2c-e**). One subset of iGluSnFR mice (n= 5) significantly increased activity at reward, t(4) = 3.75, p < 0.01. A different subset of iGluSnFR mice (n = 7) significantly decreased activity at reward, t(6)≥ = −6.43, p < 0.001. We found no systematic differences in fiber placement between reward-increasing or reward-decreasing iGluSnFR mice; implants spanned the entire VTA from anterior to posterior and subdivision (rostral linear, caudal linear, interfascicular, parabrachial pigmented, and paranigral nuclei, **Supplemental Fig. 1**). The divergent reward-related iGluSnFR activity profiles were also not explained by differences in conditioning. iGluSnFR reward groups discriminated between cues equivalently and received comparable training (**Supplemental Fig. 2**).

**Figure 2.**
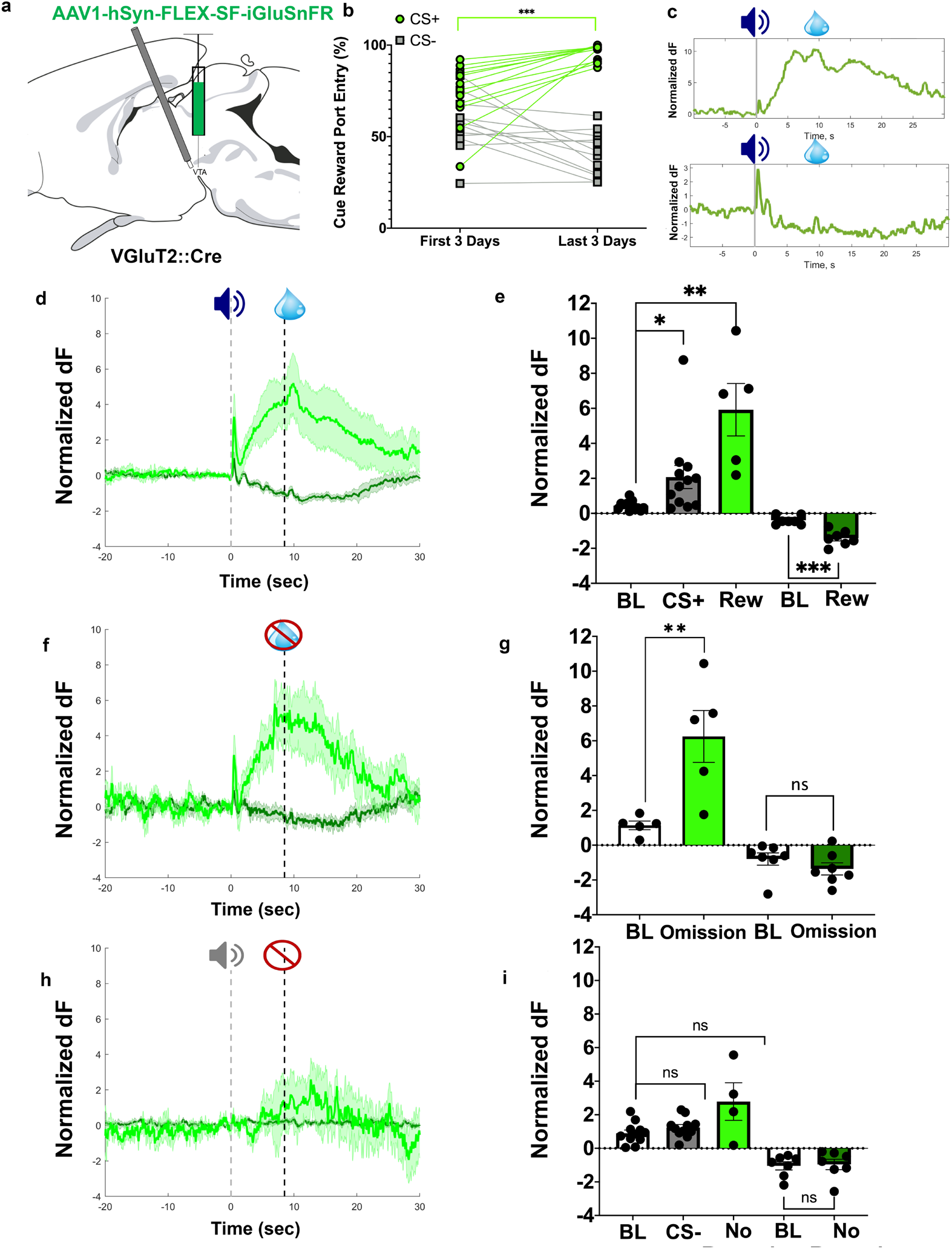
Glutamate Input (iGluSnFR) to VTA VGluT2 neurons during Pavlovian Reward. a) VGluT2::Cre mice were injected in the VTA with an AAV encoding Cre-dependent iGluSnFR and an optic was implanted in the VTA. b) Mice learned to discriminate between reward-predictive cues (CS+, green) and neutral cues (CS-, grey). c) Top: Sample trace from representative mouse with an increase at CS+ and an increase at reward (“reward-increasing” iGluSnFR mouse, n = 5). Bottom: Sample trace from representative mouse with an increase at CS+ but a decrease at reward (“reward-decreasing” iGluSnFR mouse, n = 7). d) Normalized dF of iGluSnFR subgroups during reward trials (group mean: solid line, S.E.M: shading). Reward-decreasing iGluSnFR activity increases at CS+ and decreases at reward (dark green). Reward-increasing iGluSnFR activity increases following at CS+ and reward (light green). e) Normalized dF of CS+ and reward compared to baseline. Dark green - reward decreasing, Light green - reward increasing. f) Normalized dF of iGluSnFR subgroups for reward error trials (group mean: solid line, S.E.M: shading). Dark green - reward decreasing, Light green - reward increasing. g) Normalized dF of reward omission compared to baseline. Dark green - reward decreasing, Light green - reward increasing. h) Normalized dF of iGluSnFR subgroups for neutral reward trials (CS-; group mean: solid line, S.E.M: shading). Dark green - reward decreasing, Light green - reward increasing. i) Normalized dF of CS- and non-reward outcome compared to baseline. Dark green - reward decreasing, Light green - reward increasing. * p < 0.05, ** p < 0.01, *** p < 0.001.

Mice with iGluSnFR decreases at reward did not show a significant change in neuronal activity from baseline during reward omission, t(6) = 1.347, p = 0.23. However, mice with iGluSnFR increases at reward significantly increased activity at reward omission, t(4) = −5.36, p < 0.01 (**Fig. 2f-g**). This suggests that reward-increasing glutamate inputs to VTA VGluT2 neurons signals reward anticipation and is not sensitive to expected reward omission. Both groups of iGluSnFR mice showed no significant change in neuronal activity for neutral RewCS- trials, increasing: F(2, 8) = 3.50, p = 0.08, decreasing: F(2,12) = 1.54, p = 0.25 (**Fig. 2h-i**).

### Pavlovian Aversion Task

Following reward recordings, mice were trained on a Pavlovian foot shock paradigm in a separate context where a conditioned stimulus (ShkCS+) predicted foot shock and a neutral cue (ShkCS-) predicted no shock (**Fig. 3a**). We found that this behavioral procedure resulted in learned freezing behavior, training x cue interaction F(1,6) = 22.34, p < 0.01, that was increased in response to the ShkCS+, p < 0.001, and the ShkCS-, p < 0.01. However, after conditioning time spent freezing was significantly larger in response to the ShkCS+ compared with the ShkCS-, p < 0.01, indicating that mice learned to produce more extensive fear behavior in response to the shock predicting cue (**Fig. 3b**). All mice within each sensor group received equivalent conditioning and were recorded fiber photometrically.

**Figure 3.**
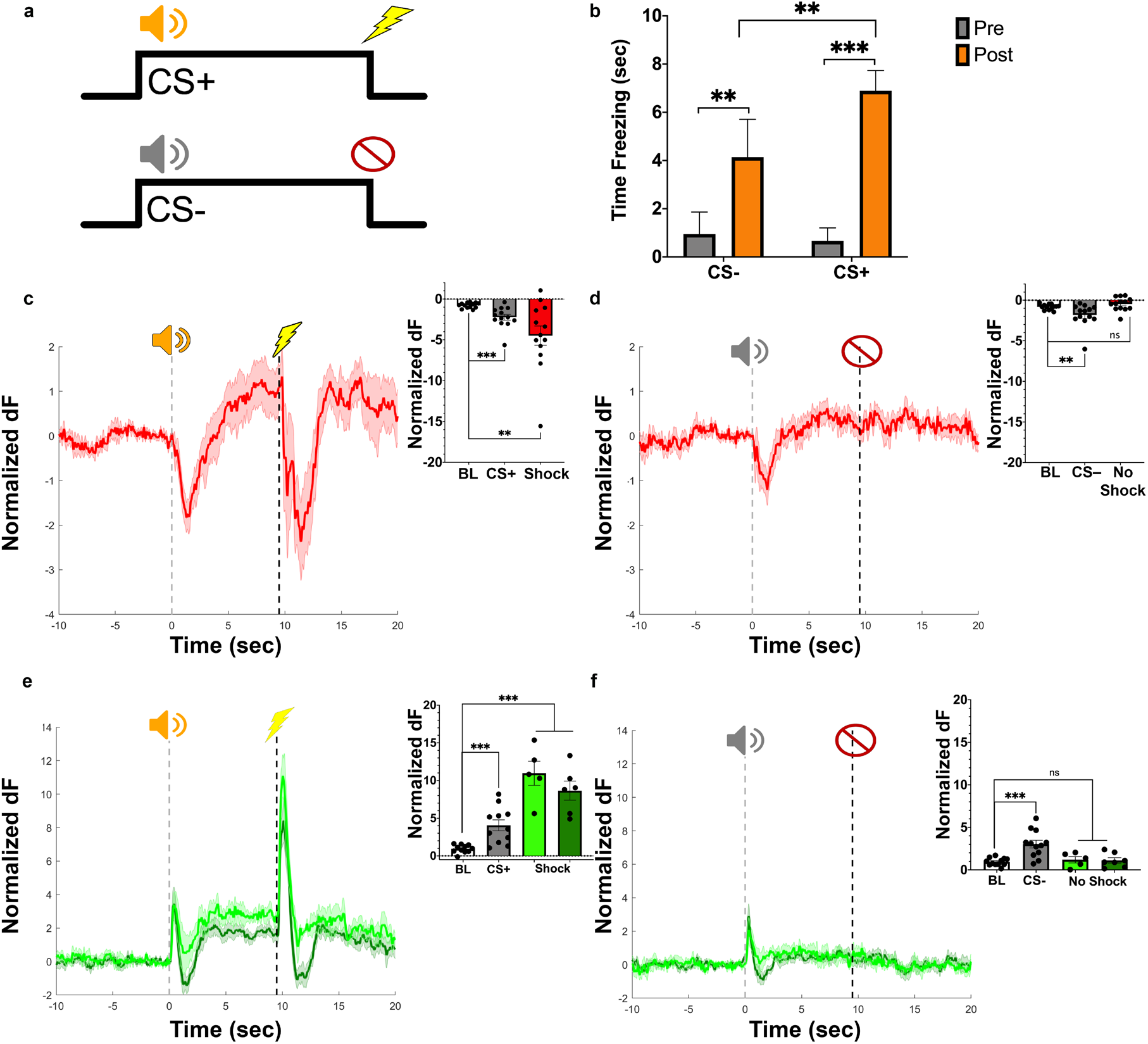
GABA and Glutamate input to VTA VGluT2 neurons during Pavlovian Aversion. a) Pavlovian aversion task. During training grid shock was paired with a CS+ tone and no shock was paired with a CS-. b) Time freezing for each cue pre-training or post-training (3 days). Training increased freezing to each cue. Freezing was significantly higher following the CS+ compared to the CS-. c) Normalized iGABASnFR dF for CS+ trials (group mean: solid line, S.E.M: shading). Inset: Significant decreases in iGABASnFR activity at CS+ and shock compared to baseline. d) Normalized iGABASnFR dF for neutral trials (CS-) (group mean: solid line, S.E.M: shading). Inset: Significant decrease in iGABASnFR activity at CS- but not non-shock outcome. e) Normalized iGluSnFR dF for CS+ shock trials (group mean: solid line, S.E.M: shading; Reward-increasing subgroup: light green; Reward-decreasing subgroup: dark green). Inset: Significant increase in iGluSnFR activity at CS+ and shock compared to baseline. f) Normalized iGluSnFR dF for neutral trials (CS-; group mean: solid line, S.E.M: shading; Reward-increasing subgroup: light green; Reward-decreasing subgroup: dark green). Inset: Significant increase in iGluSnFR activity at CS- but no change for non-shock outcome. ** p < 0.01, *** p < 0.001.

### GABA inputs to VTA VGluT2 Neurons during Pavlovian Aversion

iGABASnFR activity was modulated by aversive stimuli trials, F(2,24) = 6.48, p < 0.05. iGABASnFR activity rapidly decreased following the ShkCS+ cue, F(1,12) = 20.75, p < 0.001, and foot shock delivery, F(1,12) = 9.96, p < 0.01 (**Fig. 3c**). Additionally, iGABASnFR activity was modulated by trials predicting the absence of footshock, F(2,24) = 7.47, p < 0.01. iGABASnFR activity decreased following ShkCS-presentation, F(1,12) = 7.10, p < 0.05, but quickly returned to values not statistically different from baseline at the non-shock outcome, F(1,12) = 2.73, p = 0.12 (**Fig. 3d**).

### Glutamate inputs to VTA VGluT2 Neurons During Pavlovian Aversion

In contrast to the diverse responses of iGluSnFR activity to reward, iGluSnFR activity homogeneously increased in response to Pavlovian cues associated with footshock and footshock delivery in each iGluSnFR mouse. Increases in iGluSnFR activity did not statistically differ between glutamate sensor groups (reward increasing or reward decreasing) at any event (ShkCS+: t(10) = 0.029, p = 0.98; footshock: t(10) = −1.69, p = 0.12; ShkCS-: t(10) = 0.085, p = 0.93; non-shock outcome: t(10) = −0.17, p = 0.86). Therefore, iGluSnFR subgroup data was collapsed for within-subjects statistical comparisons. iGluSnFR activity was significantly modulated by aversive stimuli trials, F(2,22) = 65.04, p < 0.001. iGluSnFR activity significantly increased following ShkCS+ cues, F(1,11) = 23.44, p < 0.001, and following foot shock, F(1,11) = 106.21, p < 0.001 (**Fig. 3e**). iGluSnFR was also significantly modulated by trials predicting the absence of footshock, F(2,22) = 18.97, p < 0.001. While iGluSnFR activity significantly increased following neutral ShkCS-cues, F(1,11) = 23.49, p < 0.001, there was no significant difference in activity from baseline at the non-shock outcome, F(1,11) = 1.34, p = 0.27 (**Fig. 3f**).

### Neuronal activity of VTA VGluT2 Neurons During Pavlovian Reward

To identify how VTA VGluT2 neurons integrate diverse neurochemical inputs into coherent neuronal activity patterns, we recorded intracellular calcium dynamics within these neurons during the Pavlovian reward task. GCaMP mice (**Fig. 4a**) learned to emit head entries into the reward port over training depending on the cue presented, F(1,12) = 35.31, p < 0.0001. Reward port head entries for RewCS+ trials were significantly increased between the first and last three days of behavioral training, p < 0.0001, indicating that mice learned to increase responding for reward trials (**Fig. 4b**). RewCS-head entries did not change over training, p = 0.56. GCaMP activity was modulated by RewCS+ trials, F(2,12) = 11.35, p < 0.001. VTA VGluT2 neuron activity was robustly increased following the presentation of the RewCS+, F(1,6) = 28.63, p < 0.01, and remained increased at reward delivery, F(1,6) = 16.59, p < 0.001 (**Fig. 4c-4d**). Neuronal activity was also modulated by reward omission trials, F(1,6) = 9.69, p < 0.01 (**Fig 4e**). While GCaMP activity was significantly elevated over baseline at the time of reward omission, F(1,6) = 13.2, p < 0.01, we observed that the reward-related increase in GCaMP activity following reward presentation was not observed following reward omission. Direct comparison showed that reward omission activity was significantly decreased compared with reward presentation, t(6)= −2.73, p < 0.05 (**Fig. 4g-h**). The discrepancy between GCaMP activity when reward was received versus when an expected reward was omitted suggests that VTA VGluT2 neurons discriminate anticipated from received reward. Calcium signaling was not altered by RewCS- trials, F(2,12) = 0.19, p = 0.83 (**Fig. 4f, Fig. 4i**).

**Figure 4.**
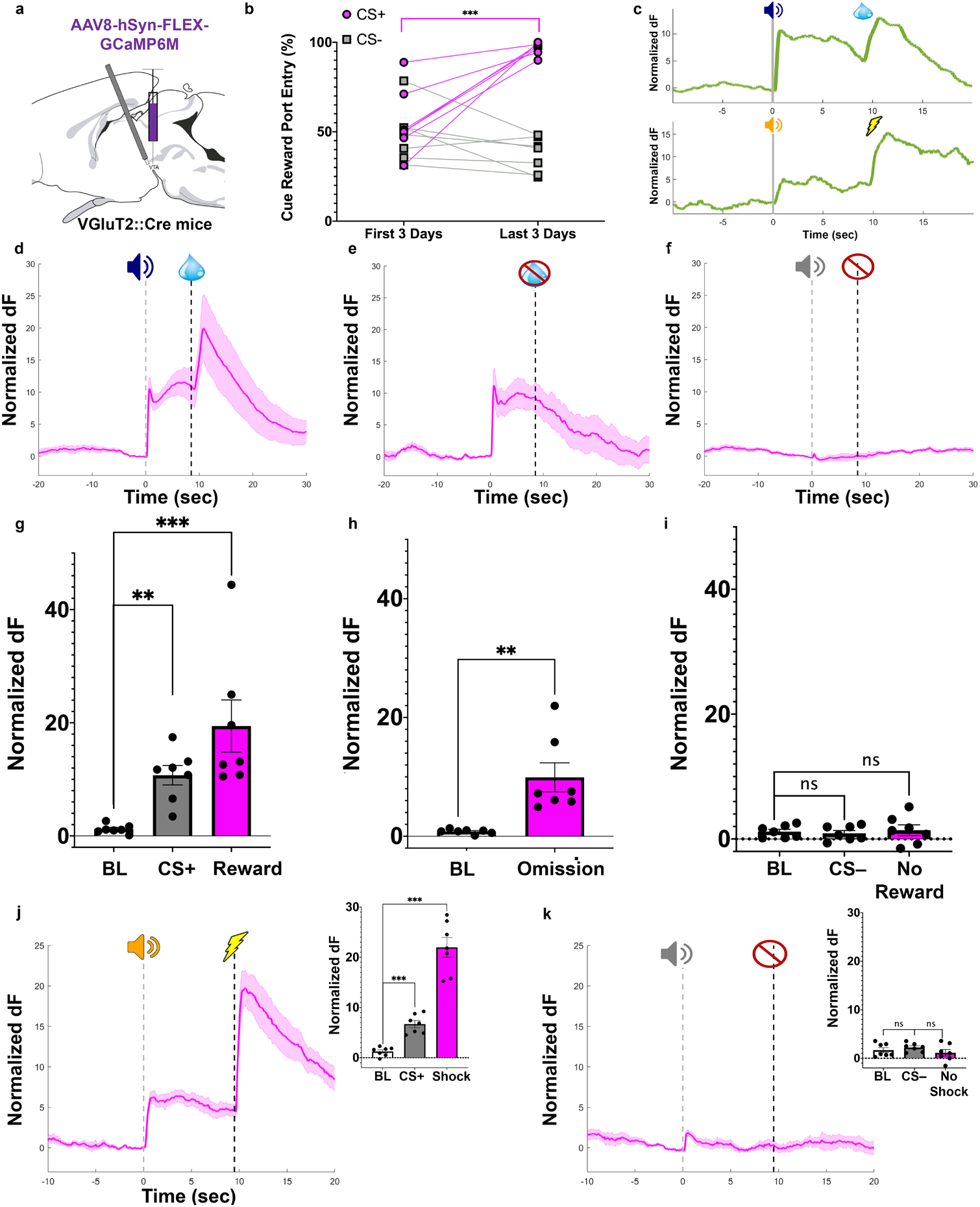
Neuronal Activity of VTA VGluT2 neurons during Pavlovian Reward and Aversion. a) VGluT2::Cre mice were injected in the VTA with an AAV encoding Cre-dependent GCaMP6m and an optic fiber was implanted in the VTA. b) Mice learned to discriminate between reward-predictive cues (CS+, purple) and neutral cues (CS-, grey). c) Top: Sample trace for GCaMP recording during Pavlovian reward, increase calcium signaling at CS+ and sucrose reward. Bottom: Sample trace for GCaMP recording during Pavlovian aversion, increase at shock CS+ and shock. d-f) Normalized dF for reward trials (d), reward error trials (e), and neutral CS- trials (f) (group mean: solid line, S.E.M: shading). g-i) Quantification for d-f. Comparison of baseline GCaMP activity for CS+ and reward (g), reward omission (h), and CS- and non-shock outcome (i). Significant increases in GCaMP activity at CS+, CS-, reward and reward omission. No significant changes for neutral CS- trials. j) Normalized GCaMP dF for shock trials (group mean: solid line, S.E.M: shading). *Inset*: Significant increase in GCaMP activity at CS+ and shock compared to baseline. k) Normalized GCaMP dF for neutral trials (CS-; group mean: solid line, S.E.M: shading). *Inset*: No significant change in GCaMP activity during CS- trials compared to baseline. ** p < 0.01, *** p < 0.001.

### Neuronal activity of VTA VGluT2 Neurons During Pavlovian Aversion

Given that iGluSnFR activity increased and iGABASnFR activity decreased during shock, we hypothesized that VTA VGluT2 neuronal activity would increase by an aversive outcome. VTA VGluT2 neuron activity was modulated by shock trials, F(2,12) = 60.047, p < 0.001. GCaMP activity significantly increased following presentation of the ShkCS+, F(1,6) = 44.74, p < 0.001, as well as foot shock, F(1,6) = 80.44, p < 0.001 (**Fig. 4j**). Calcium signals were not significantly altered by ShkCS- trials, F(2,12) = 1.39, p = 0.29 (**Fig. 4k**).

### Changes in firing of VTA VGluT2 Neurons by Glutamate and GABA Receptor Modulation

Our results indicate that glutamate and GABA inputs to VTA VGluT2 neurons are dynamically regulated by reward and aversion-based learned cues and outcomes. However, it is unclear how glutamate and GABA affect the firing of VTA VGluT2 neurons. To determine how glutamate and GABA affects the firing of VTA VGluT2 neurons, we fluorescently labeled VTA VGluT2 neurons by intra-VTA injection of Cre-dependent eYFP in VGluT2::Cre mice and performed cell-attached and whole-cell electrophysiological recordings of eYFP-expressing neurons in VTA-containing slices (**Fig. 5a**). Glutamate and/or GABA receptor antagonists or agonists were washed onto slices and spontaneous or evoked firing of VTA VGluT2 neurons was assessed. VTA VGluT2 neuron firing was significantly decreased following application of the AMPA and NMDA antagonists DNQX/APV, t(5) = 5.64, p < 0.01, indicating that glutamate input maintains VTA VGluT2 basal firing (**Fig. 5b**). Surprisingly, when the GABA-A receptor antagonist Bicuculine was applied, VTA VGluT2 neurons significantly decreased firing rate t(5) = 2.72, p < 0.05 (**Fig. 5c**). This result was contrary to our initial hypothesis that VTA VGluT2 neurons would be disinhibited by GABA-A receptor blockade. Further, co-application of bicuculline and DNQX/APV decreased excitability of VTA VGluT2 neurons as measured by evoked spiking in response to current injection. We observed a significant increase in the number of action potentials with increasing current injection, F(9,72) = 63.28, p < 0.0001, and a significant effect of treatment, F(1, 8) = 5.94, p < 0.05, indicating that GABA-A receptor blockade does not rescue decreased firing induced by AMPA/NMDA receptor blockade (**Fig. 5 d-e**). VTA VGluT2 neuron firing was significantly increased following application of AMPA/NMDA, t(5) = 3.461, p = 0.018, but did not significantly change firing following application of the GABA-A receptor agonist Isoguvacine, t(5) = 1.19, p = 0.11 (**Fig. 5 f-g**). These results indicate that glutamate is the primary driver of increasing or decreasing neuronal activity of VTA VGluT2 neurons. In contrast, GABA inputs acting at GABA-A receptors are surprisingly ineffective at changing VTA VGluT2 neuron basal firing rates.

**Figure 5.**
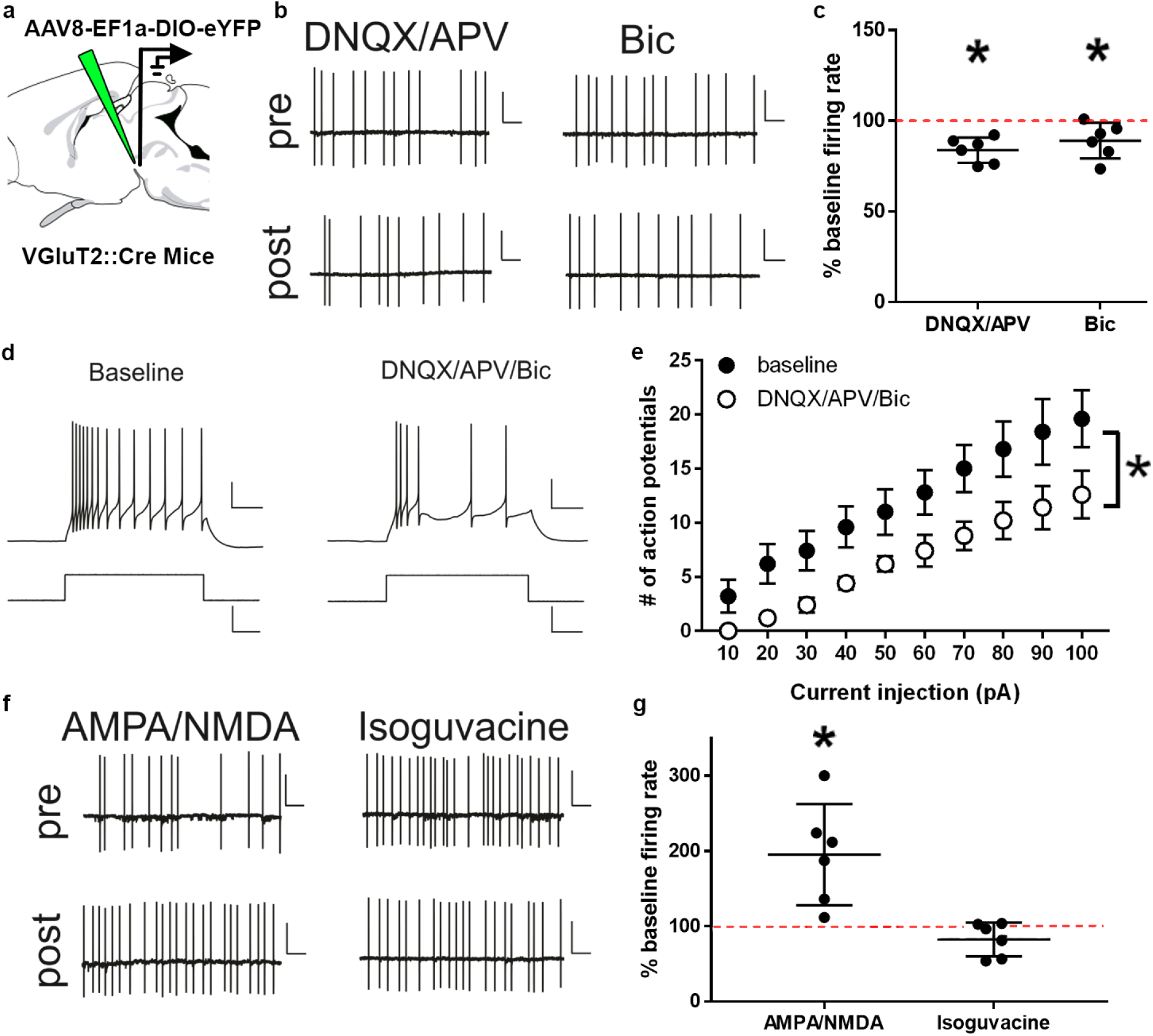
VTA VGluT2 neuron firing following GABA/Glutamate receptor agonism and antagonism. a) VGluT2::Cre mice were injected in the VTA with an AAV encoding Cre-de- pendent enhanced yellow fluorescent protein (eYFP). eYFP-expressing cells were recorded ex vivo in whole-cell or cell-attached mode. b-c) Both AMPA/NMDA (DNQX/APV) and GABA-A (bicuculline) receptor antagonists reduced baseline firing in VTA VGluT2 neurons. c) Quantification for each recorded cell following DNQX/APV or bicuculline compared to baseline firing rate. d) Representative firing from a VTA VGluT2 neuron following AMPA/NMDA/GABA-A receptor antagonist combination following current injection. e) Excitability of VTA VGluT2 neurons was reduced compared to baseline following combined AMPA/NMDA/GABA-A receptor antagonism. f-g) Glutamate receptor agonists (AMPA/ NMDA) increase firing in VTA VGluT2 neurons but GABA-A receptor agonists (isoguvacine) had no significant effect. g) Quantification for each recorded cell following AMPA/ NMDA or isoguvacine compared to baseline firing. * p < 0.05

## Discussion

Slightly over a decade ago the cellular landscape of the VTA expanded to include populations of neurons that express VGluT2, a molecular marker for neurons that accumulate glutamate into synaptic vesicles^26^. Since their discovery, subpopulations of VGluT2 neurons have been categorically divided based on their firing for rewarding and aversive stimuli^16^, projection targets^27^, and neurotransmitter co-release characteristics^19,28–30^. Thus far, VTA VGluT2 neurons appear to have diverse roles in motivated behavior^25,26^. Here, by using genetically encoded fluorescent reporters of glutamate and GABA signaling^31–33^, we aimed to identify how diverse rewarding and aversive information is neurochemically communicated to VTA VGluT2 neurons during motivated behavior.

We found that the pattern of glutamate inputs to VTA VGluT2 neurons during reward is not homogenous. Specifically, one subpopulation of glutamate inputs increased while another decreased at reward delivery. Electrophysiological recordings from optogenetically-identified VTA VGluT2 neurons have also identified subtypes of neurons based on whether they increase or decrease firing rate at the presentation of reward^16^. Importantly, differences in recorded glutamate input at reward was not systematically explained by topographical fiber/viral expression location within the VTA or behavioral training. Our recordings with iGluSnFR within VTA VGluT2 neurons suggest that distinct populations of VTA VGluT2 neurons receive quantitatively heightened or reduced glutamate when an appetitive reward is presented.

We interpret the differences in recorded glutamate input as reflecting distinct glutamate projections to VTA VGluT2 neurons that at least partially mediate the diverse firing patterns of VTA VGluT2 neurons in response to reward. If this is the case, then it would be expected that increased glutamate input would correspond with enhanced firing and decreased glutamate input with a reduction in firing. Indeed, we observed that activating glutamate receptors was sufficient to increase VTA VGluT2 neuron firing and blocking glutamate receptors was sufficient to decrease VTA VGluT2 neuron firing ex vivo. We hypothesize that the functional heterogeneity of glutamate inputs to VTA VGluT2 neurons arises from circuit-level differences within this subset of neurons. Further research will be necessary to test whether distinct populations of VTA VGluT2 neurons receive functionally separate glutamate inputs from unique brain regions^34^, target different output brain regions^27^, or belong to a population of genetically unique VTA VGluT2 neurons that releases one or more neurotransmitters^29,30,35^.

In contrast to glutamate inputs, GABAergic inputs to VTA VGluT2 neurons, measured by iGABASnFR, showed homogeneously decreased activity in response to reward. Our *ex vivo* slice recordings indicated that although these neurons receive GABAergic input, VTA VGluT2 neuronal firing is more potently modulated by glutamate inputs than GABA inputs. To assess how VTA VGluT2 neurons integrate glutamate, GABA, and other inputs into a coherent neuronal activity pattern, we performed GCaMP recordings of VTA VGluT2 neurons in the same behavioral paradigm. Despite a heterogeneity in reported firing within these cells at reward^16^, and our demonstration of diverse patterns of glutamate inputs to these cells at reward, GCaMP recordings showed homogenously increased activity during reward. It is possible that integrating glutamate and GABA input at the reward event requires increased calcium signaling. Alternatively, because our GCaMP recordings were measured fiber photometrically rather than at a single-neuron level, it is possible that we were unable to resolve the diversity of VTA VGluT2 neuron responses to reward using GCaMP.

Despite their diverse signaling of reward, glutamate inputs to VTA VGluT2 neurons homogenously increased by aversive cues and events. GABAergic input to these neurons decreased following aversive cues and events. Thus, we interpret the enhanced glutamatergic input to VTA VGluT2 neurons, as well as decreased GABA input in response to aversive stimuli, as a mechanism that increases the activity of VTA VGluT2 neurons in response to aversive stimuli. Indeed, most VTA VGluT2 neurons increase firing rates in response to aversive facial airpuffs^16^.

A subset of VTA VGluT2 neurons increases firing in response to both rewarding and aversive stimuli, suggesting this subpopulation may function as salience detectors^16^. A separate subset of VTA VGluT2 neurons decreases firing in response to reward and increases firing to aversive stimuli, suggesting they may participate in aversion. Our iGluSnFR recordings mirror these results, identifying one population of glutamate inputs that increases in response to both rewarding and aversive stimuli, and another population of glutamate inputs that decreases in response to reward but increases in response to aversive stimuli. Together, we hypothesize that distinct glutamatergic inputs to VTA VGluT2 neurons drive their signaling of and participation in differently valenced behaviors that have been observed following their optogenetic activation^17–20^.

Given that VTA dopamine neurons signal negative prediction error following unexpected reward omission^36^, we aimed to identify if glutamate or GABA inputs to VTA VGluT2 neurons are sensitive to negative prediction errors. The pattern of activity for glutamate and GABA inputs to VTA VGluT2 neurons was not different between reward trials and reward error trials, suggesting that these neurotransmitter inputs signal anticipation of an expected reward but are insensitive to whether the outcome is received. Interestingly, when recording VTA VGluT2 intracellular calcium signaling via GCaMP, we found that the activity of VTA VGluT2 neurons was reduced following reward omission compared with reward receipt. This suggests that VTA VGluT2 neurons process the omission of an expected reward but that their GABAergic or glutamatergic inputs are not responsible for relaying this information. These data highlight the unique opportunities that cell-type specific neurotransmitter sensors offer when used in combination with neuronal activity sensors such as GCaMP in identifying how specific neurochemicals influence motivation-related neuronal activity. Further research is necessary to identify which neurochemical inputs to VTA VGluT2 neurons distinguish reward from unexpected reward omission. As some VTA VGluT2 neurons ex-press D2 receptors, one candidate is dopamine neuron input^37^.

Glutamate and GABA inputs to VTA VGluT2 neurons were modulated by learned predictors (conditioned stimuli) of rewarding and aversive outcomes. Glutamate input to VTA VGluT2 neurons was significantly increased at the presentation of reward-predicting (RewCS+) and aversion-predicting (ShkCS+) cues. Previous work established that VTA VGluT2 neurons increase firing following aversive air puff but this is the first identification, to our knowledge, that these neurons signal the anticipation of aversion. Our iGluSnFR data in coordination with GCaMP recordings, demonstrate that VTA VGluT2 neurons are indeed activated by cues predicting aversive outcomes involving at least direct glutamatergic excitation. Altogether, it appears that select glutamate inputs to VTA VGluT2 neurons may more reliably signal reward-related events than others, but glutamate inputs are strongly relayed to VTA VGluT2 neurons following aversion-related cues and events. GABA inputs on the other hand, decrease activity following reward or aversion-related predictors and subsequent outcomes. During reward conditioning, cues that predicted the absence of sucrose delivery did not alter neurotransmitter or calcium signaling to VTA VGluT2 neurons. While no sensor detected a significant change in activity following the RewCS-, all glutamate inputs significantly increased and GABA inputs significantly decreased following the ShkCS-that predicted the absence of footshock. It is possible that the ShkCS-signaling reflected a generalized freezing behavior to this stimulus, though significantly less than the ShkCS+. Alternatively, it is also the possible the ShkCS-served a “safety” signal that might be interpreted as reward-predicting^38,39,40^.

We examined whether the electrophysiological diversity of VTA VGluT2 is related to increasing or decreasing glutamatergic excitation or GABAergic disinhibition. To identify how glutamate and GABA affect VTA VGluT2 neuron firing, we recorded from VTA VGluT2 neurons ex vivo in the presence or absence of glutamate and/or GABA receptor antagonists or agonists. Both AMPA/NMDA and GABA-A receptor antagonists significantly reduced VTA VGluT2 neuron firing compared to baseline. The decreased firing following AMPA/NMDA receptor antagonism indicates that tonic level of glutamate input is necessary for maintaining baseline firing rates in VTA VGluT2 neurons. However, contrary to our hypothesis GABA-A receptor antagonism reduced VTA VGluT2 neuron firing instead of increasing it. It is unlikely that the concentration of bicuculline was too low to detect changes in firing because the concentration used is sufficient to disinhibit VTA dopamine neurons by nearly 50% of basal rates^41^. Instead, this result might be explained by the expression of GABA-A receptors on GABAergic inputs that gate glutamate inputs to these neurons. Regardless of the mechanism, this result indicates that a reduction in GABA-A receptor activity is insufficient to increase VTA VGluT2 neuron firing through a disinhibitory mechanism. Similar results were found from experiments using AMPA/NMDA and GABA-A receptor agonists. VTA VGluT2 neurons robustly increased spontaneous firing following AMPA/NMDA application but were unaffected by GABA-A receptor agonism. In addition, combining AMPA/NMDA antagonists with a GABA-A receptor antagonist did not rescue the decreased firing rates. It was again surprising that GABA-A receptor agonism did not reduce VTA VGluT2 neuron firing, however this may be because VGluT2 neurons rest relatively close to the reversal potential for chloride^37^ leading to little ion flux through GABA-A channels at resting potentials. It is unlikely that the concentration of isoguvacine was too low to detect changes in firing because the concentration used is sufficient to reduce VTA dopamine neuron firing by nearly 50% of basal rates^42^. Taken together, our electrophysiological results indicate that changes in firing by VTA VGluT2 neurons are driven by increasing or decreasing fluctuations in glutamate input and are remarkably less influenced by GABA-A receptor mediated GABA transmission. Specifically, removal of glutamate receptor activity results in disexcitation of VTA VGluT2 neurons while activation of glutamate receptor activity increases VTA VGluT2 neuron firing. While GABA-A receptors do not contribute substantially to basal firing, it is likely they still play an important role in regulating VTA VGluT2 neurons, perhaps through regulating integration of excitatory inputs. It is also plausible that GABA-B receptors or other inhibitory neurotransmitters such as glycine play a larger role in inhibiting VTA VGluT2 neurons.

VTA VGluT2 neurons process and signal multiple aspects of motivated behavior, including reward and aversion^16–19^. Here we identified that glutamate and GABA inputs to VTA VGluT2 neurons are modulated by reward and aversion predicting cues and outcomes. Glutamate inputs in particular, show the same diverse responses of VTA VGluT2 neurons following reward as those measured by electrophysiological firing patterns^16^. Together with the identification that glutamate receptor activity principally regulates VTA VGluT2 neuron firing, our results suggest that distinct glutamate inputs drive VTA VGluT2 neuron participation in different motivated behaviors. Lastly, our results indicate that neurotransmitter sensors serve as a powerful tool to identify neurochemical influences that control motivated behavior and expand on the electrophysiological firing patterns that underlie them.

## Acknowledgments

Research reported in this publication was supported by the National Institute On Drug Abuse of the National Institutes of Health under Award Number R01DA047443 (DHR). The content is solely the responsibility of the authors and does not necessarily represent the official views of the National Institutes of Health. This research was also supported by the Webb-Waring Biomedical Research Award from the Boettcher Foundation, the CO-PILOT award from the Colorado Clinical and Translational Sciences Institute, a 2020 NARSAD Young Investigator grant from the Brain and Behavior Research Foundation, and The University of Colorado (DHR). Further support was provided by R00 MH106757 and a 2019 NARSAD Young Investigator grant (AMP). We thank Drs. Loren Looger and Jonathan Martin for generously providing viruses encoding Cre-dependent iGluSnFR and iGABASnFR, to Dr. Karl Deisseroth for generously providing the Cre-dependent eYFP virus, and to Addgene for providing the Cre-dependent GCaMP vector and Alysabeth Phillips for photometry piloting and other technical assistance. The funders had no role in study design, data collection and analysis, decision to publish, or preparation of the manuscript.

## Author contributions

DJM performed stereotactic injections, surgeries, fiber photometry recordings, behavioral training, and photometry and behavioral analysis. AMP performed slice electrophysiological recordings in VTA VGluT2 neurons and directly contributed data for Figure 5. DHR contributed behavioral and photometry data analysis, histology, and stereotactic surgeries.

## Conflict of interest statement

The authors have no financial interests to be disclosed.

---

**Supplemental Figure 1:**
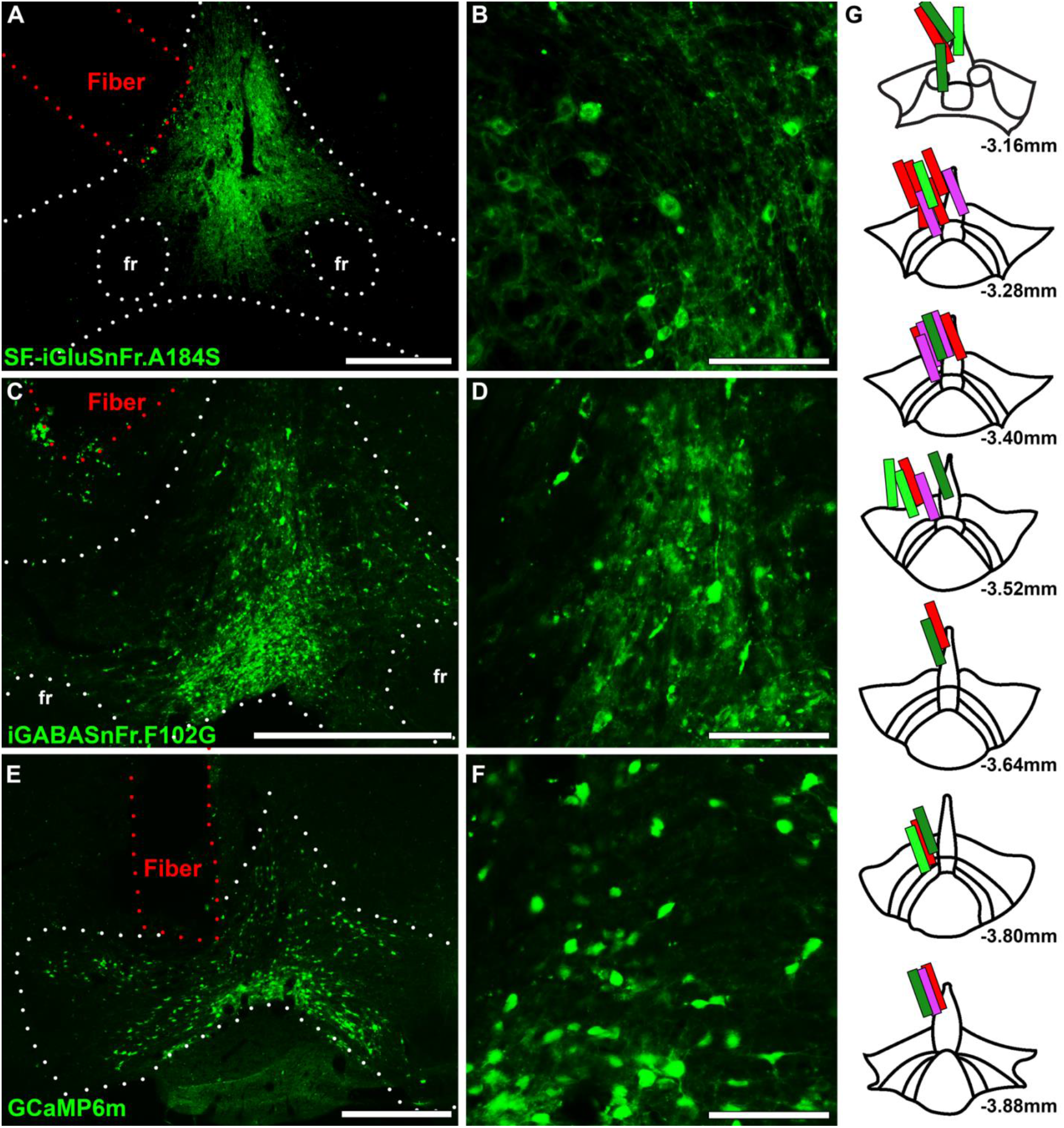
Topography of fiber implant locations for all sensors. **a-b.** Example iGluSnFR cellular expression within VTA VGluT2 neurons in low (a) and high (b) magnification. **c-d.** Example iGABASnFR cellular expression within VTA VGluT2 neurons in low (a) and high (b) magnification. **e-f**. Example GCaMP cellular expression within VTA VGluT2 neurons in low (a) and high (b) magnification **g**. Fiber localizations. Light green – iGluSnFR reward increasing subgroup; Dark green – iGluSnFR reward decreasing subgroup; Red – iGABASnFR; Purple –GCaMP. Number refers to anteroposterior position from bregma (mm). Scale bars: A,C,E – 250 μm, B,D,F – 100 μm. fr – fasciculus retroflexus.

**Supplemental Figure 2:**
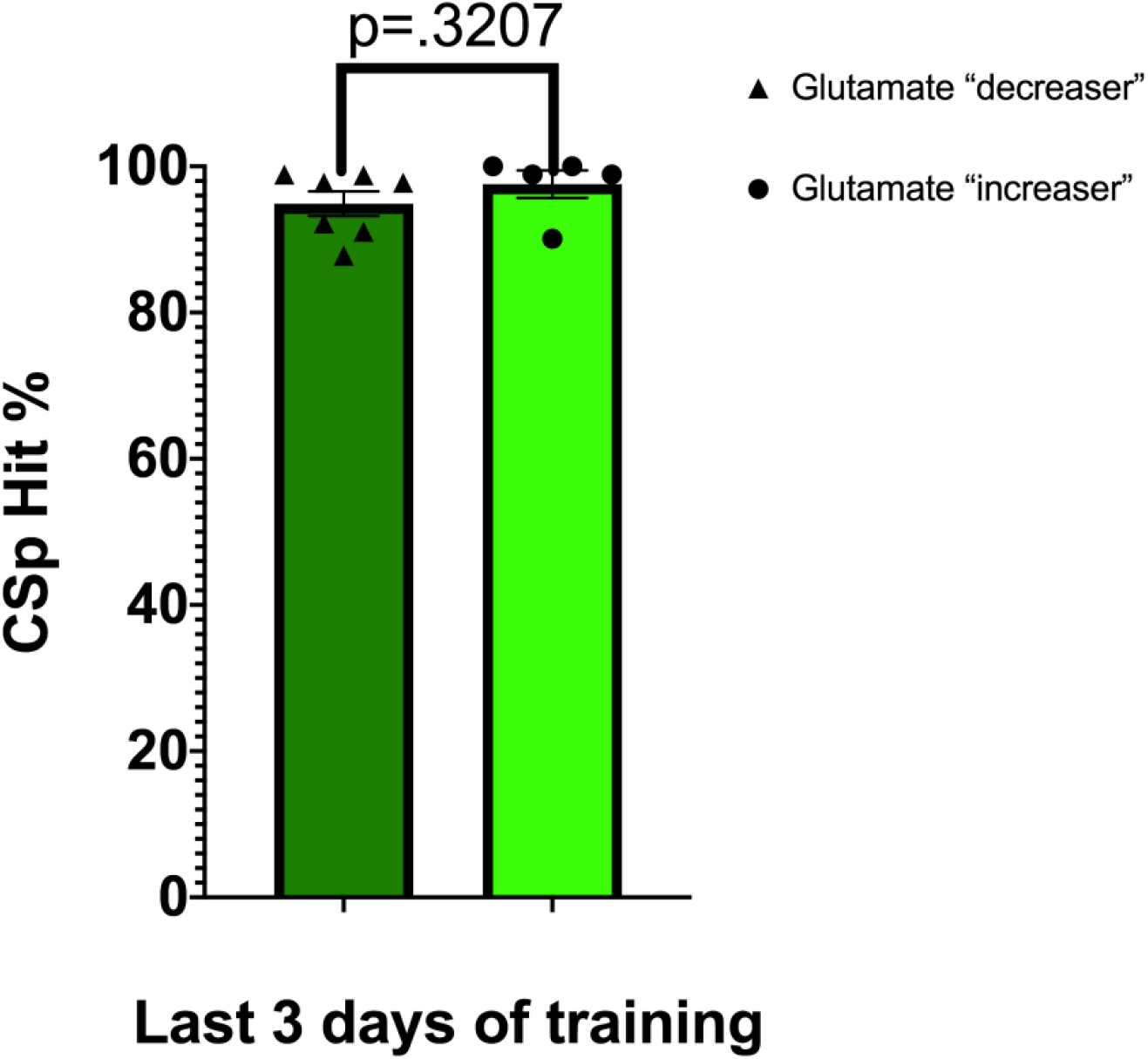
Reward training for iGluSnFR animals shows no significant behavioral difference between reward subgroups. Percent of RewCS+ trials in which mice entered the reward port for iGluSnFR reward increasing and iGluSnFR reward decreasing subgroups during the last three days of training.

## Notes

### Competing Interest Statement

The authors have declared no competing interest.

